# Ancient DNA analyses of two Early Chalcolithic Individuals from the West Mound

**DOI:** 10.1101/2024.10.22.619570

**Authors:** Ayça Doğu, Damla Kaptan, Eren Yüncü, Kanat Gürün, Kıvılcım Başak Vural, Maciej Chylenski, Jennifer Byrnes, Anders Götherström, Füsun Özer, Mehmet Somel

## Abstract

In this chapter, we investigate the genetics of two Early Chalcolithic Çatalhöyük individuals, U.18333 and U.16835, the only two burials yet recovered from the West Mound. These were two neonates buried within the same building. Using shotgun-sequenced partial genomes (0.06x and 0.02x coverage) we identify both as females. Despite being recovered from the same building, we find no close genetic kinship between them, in line with previously published kinship results from Çatalhöyük. We also find that the two West Mound neonates shared the same gene pool with Neolithic Çatalhöyük and other Central and West Anatolian Neolithic populations, and they did not carry the Caucasus-related “eastern” gene flow signature observed in later-coming Chalcolithic Anatolian genomes. This indicates no large-scale admixture between East and West Mound Çatalhöyük, and possibly that the post-Neolithic “eastern” gene flow event into Anatolia may have initiated only by the mid-6th millennium BCE.

## Introduction

Here we present partial genomes from the only two individuals yet excavated at Çatalhöyük West Mound. These are two neonates, U.18635 and U.18333, buried in building B.105 (Byrnes and Anvari, this volume), and have been dated to 6000-5800 cal. BC (95%) and 5900-5700 cal. BC (95%) based on SUERC-100610 and SUERC-100611, respectively, using a Bayesian stratigraphic model (Rosenstock et al., this volume). With the available data, we address two questions: whether the two represented members of the same biological family, and whether they carry genetic signatures of inferred population movements in the Early Chalcolithic period.

Regarding the first question, bioarchaeological and genetic studies at Çatalhöyük, as well as at other Neolithic Anatolian sites, have been reporting intriguing patterns of genetic kinship among co-buried individuals, i.e. in-site burials within the same spaces. On the one hand, published data from 9th and 8th millennium Aceramic/Pre-Pottery Neolithic (PPN) sites, Aşıklı Höyük and Boncuklu of Central Anatolia (Yaka, Mapelli, et al. 2021), and Çayönü of the Upper Tigris Basin (Altınışık et al. 2022) suggest that co-burials within buildings were frequently close biological kin, including first-, second- (e.g. avuncular) and third-degree (e.g. first cousins) relationships. On the other hand, data from the 7th millennium BCE sites of Çatalhöyük East Mound and the Northwest Anatolian Barcın Höyük suggest a lower frequency of close biological kin among co-buried individuals. The Çatalhöyük evidence includes dental non-metric traits of 266 individuals, mainly of adults, showing no indication of within-building clustering compared to contemporaneous buildings (Pilloud and Larsen 2011), as well as high mitochondrial haplogroup diversity identified across 10 adults and subadults buried in three buildings (Chyleński et al. 2019). More recently, Yaka and colleagues used partial genomes to estimate biological kinship levels among nine subadults buried in three buildings and could only identify a single pair of close biological kin in this sample; the data from Barcın was of similar nature (Yaka, Mapelli, et al. 2021). Overall, these results imply that in Çatalhöyük and in Barcın, in apparent contrast to Aceramic/PPN sites, social organization at the level of burial location choices may not have been focused on biological kinship. Multiple biological families appear to have used the same spaces to bury their dead, including newborns and children [for discussion on the implications of such traditions, see e.g. (Larsen et al. 2015) and (Hodder 2022)]. It is therefore interesting to investigate whether this tradition may have continued into the 6th millennium BCE, even though our sample is limited to two West Mound neonates.

The second question we address is on population genetics of early Holocene Anatolia; specifically, whether these two individuals carried signatures of the eastern gene flow events identified during the Chalcolithic period in Anatolia. Archaeogenomic evidence suggests that starting with the 6^th^ millennium BCE, there occurred gene flow between populations in Anatolia and groups to its east (Altınışık et al. 2022; Harney et al. 2018; Lazaridis et al. 2016, 2017, 2022; Skourtanioti et al. 2020; Wang et al. 2019). These “eastern” populations include the Caucasus, Upper Mesopotamia, or the Zagros, all of which share genetic profiles that can be distinguished from those in Central Anatolia. The 6^th^ millennium BCE population movements from Anatolia eastward are inferred from Central Anatolian Neolithic-related components arising in 6^th^ millennium BCE Caucasus and Zagros genomes (Lazaridis et al. 2016, 2022; Wang et al. 2019) (Koptekin et al. under revision). Meanwhile, a movement from the east into Anatolia can also be clearly discerned. Novel “eastern” genetic components, which could be representing Caucasus, Mesopotamia, and/or Zagros populations, appear in published post-Neolithic Central and West Anatolian genomes from the mid-6th to 4th millennia BCE, such as Büyükkaya (Central Anatolia) and Kumtepe (West Anatolia) (Altınışık et al. 2022; Kılınç et al. 2016; Skourtanioti et al. 2020), as well as 4th millennium sites such as Çamlıbel Tarlası (Central Anatolia) and İkiztepe (North Anatolia) (Lazaridis et al. 2022) (Koptekin et al. under revision). This represents a significant change in the Anatolian gene pool, with roughly one third of the post-Neolithic Anatolian ancestry assigned to early Holocene eastern sources (i.e. east of Central Anatolia) (Lazaridis et al. 2022) (Koptekin et al. under revision).

The timing, tempo (gradual vs. episodic) and the origins (the source populations) of this eastern gene flow event are yet largely unclear, although recent work has started to provide some insights. For instance, Altınışık and colleagues (2022) have estimated that early Holocene Caucasus (the so-called Caucasus hunter-gatherer genomes) are a more likely source of this post-Neolithic gene flow than Upper Mesopotamia (represented by Çayönü). Meanwhile, using the distribution of admixture tracks in post-Neolithic Anatolian genomes, Skourtanioti and colleagues (2020) estimated this eastern admixture event to have occurred around c.6500 BCE. This estimate, however, should be taken with caution, as the available genomic data from the 7th millennium BCE do not reveal any major change in West or Central Anatolian gene pool during this period (Yaka, Mapelli, et al. 2021) (Koptekin et al. under revision).

Genomic data from the two West Mound neonates could provide important insights into when such change may have happened and/or whether it impacted the Çatalhöyük population. It is particularly attractive to ask whether the shift in lifestyles between the East and West Mound (Brady et al. 2022) (#Biehl et al. summary#) may be somehow related to this inferred admixture event.

### Experimental procedures and data pre-processing

We followed standard ancient DNA isolation and sequencing library preparation procedures on skeletal material from the two West Mound neonates, U.18333 and U.16835. See (Yaka, Mapelli, et al. 2021) for details, and (Yaka, Doğu, et al. 2021) for an overview of the ancient genomics pipeline. Briefly, for each individual, we cut and ground the otic capsule part of the petrous bone, dissolved the powder and isolated DNA following (Dabney et al. 2013) (without applying UDG treatment), and prepared sequencing libraries according to (Meyer and Kircher 2010). After amplification, purification and quantification, we shotgun sequenced the libraries on the Illumina Novaseq platform, producing 11,183,100 and 11,821,310 reads (2x100 cycles) from each. After removing adapter sequences in raw FASTQ files (Schubert et al. 2016), we merged paired-end reads and aligned these to the human reference genome version hs37d5 using BWA (Li and Durbin 2009) and filtered for mapping quality 30. Next, we called pseudohaploid SNPs using two SNP reference panels, the Human Origins dataset (Patterson et al. 2012) used for principal components analyses, and an outgroup-ascertained dataset (SNPs identified in 1000 Genomes Yoruba individuals from West Africa; (Auton et al. 2015)) for D-tests (Kılınç et al. 2016; Yaka, Mapelli, et al. 2021).

### Ancient genome data overview and contamination estimates

Using direct shotgun sequencing we generated sequence data for the West Mound neonates, U.18333 and U.16835. Statistics summarizing the data, along with data from 13 published partial genomes from East Mound Çatalhöyük (Yaka, Mapelli, et al. 2021) are presented in Table 1.

**Table 1.** A summary of genetic data of the Çatalhöyük West Mound (Early Chalcholithic) individuals presented here and published Çatalhöyük Neolithic individuals (Yaka et al., 2021).

| Period | Excavation ID | Sample ID | Average Read Length | Genome Coverage (X) | Mitochondrial Genome Coverages (X) | Genetic Sex | C14 Laboratory code | Calibrated C14 age BCE |
| --- | --- | --- | --- | --- | --- | --- | --- | --- |
| E. Chalcolithic | U.18333 | cch351 | 82.87 | 0.0617 | 4.26 | XX | SUERC-100611 | 5880-5725 (95%) |
| E. Chalcolithic | U.16835 | cch352 | 80.79 | 0.0222 | 2.29 | XX | SUERC-100610 | 5900-5800 (95%) |
| Neolithic | U.5357 | CCH144 | 59.19 | 0.0594 | 2.48 | XY | OxA-21263 | 7035–6680 (93%),<br>6670–6650 (2%) |
| Neolithic | U.2017 | CCH163 | 58.63 | 0.0323 | 3.31 | XX | UCIAMS-143259 | 6815–6790 (2%),<br>6775–6595 (93%) |
| Neolithic | U.21855 | CCH285 | 64.52 | 0.0738 | 2.14 | XX | - | - |
| Neolithic | U.1885 | CCH289 | 59.63 | 0.0685 | 2.54 | XY | UCIAMS-143258 | 6905–6885 (1%),<br>6825–6635 (92%),<br>6625–6600 (2%) |
| Neolithic | U.2033 | CCH290 | 69.09 | 0.0125 | 0.37 | XY | UCIAMS-206232 | 6690–6590 (95%) |
| Neolithic | U.2779 | CCH294 | 71.57 | 0.2705 | 11.26 | XY | - | - |
| Neolithic | U.8587 | CCH311 | 71.97 | 0.1375 | 8.44 | XX | - | - |
| Neolithic | U.30006 | cth006 | 59.97 | 0.0738 | 47.73 | XX | OxA-37143<br>TÜBİTAK-684 | 6645–6495 (94%),<br>6490–6480 (1%) |
| Neolithic | U.20217 | cth217 | 54.88 | 0.0549 | 6.07 | XX | TÜBİTAK-686 | 6415–6240 (95%) |
| Neolithic | U.2728 | cth728 | 57.00 | 0.0819 | 7.45 | XX | TÜBİTAK-741 | 6695–6505 (95%) |
| Neolithic | U.11739 | cth739 | 54.21 | 0.2012 | 58.12 | XX | TÜBİTAK-685 | 6235–6075 (95%) |
| Neolithic | U.5747 | cth747 | 62.15 | 0.1224 | 52.99 | XX | OxA-33198<br>TÜBİTAK-743 | 6640–6490 (95%) |
| Neolithic | U.2842 | cth842 | 54.46 | 0.0864 | 31.05 | XX | TÜBİTAK-742 | 6690–6505 (95%) |

U.18333 and U.16835 were estimated to carry 20.9% and 7.3% endogenous aDNA, which is similar to endogeneous aDNA percentages of published Neolithic Çatalhöyük genomes from subadults (Yaka, Doğu, et al. 2021; Yaka, Mapelli, et al. 2021). We obtained genome-wide coverages of 0.06- and 0.02-fold (roughly corresponding to 6% and 2% of the genome being observed), which translated into 34550 and 12676 SNPs in the Human Origins dataset, and 288854 and 105970 SNPs in the 1000 Genomes Yoruba dataset, respectively. Mitochondrial genome coverages were 4.3- and 2.3-fold, respectively, too low to reliably detect mitochondrial DNA haplogroups. We identified both neonates as genetically female using Rx and Ry methods (Mittnik et al. 2016; Skoglund et al. 2013).

We estimated potential contamination using two approaches. The 5’ and 3’ end C->T and G->A postmortem transitions occurred at rates of >0.35 for both libraries, suggesting that the bulk of the molecules were ancient (as opposed to modern-day contamination). Contamination estimated using the contamMix and Schmutzi methods based on mitochondrial haplotypes (Fu et al. 2013; Renaud et al. 2015) also suggested the lack of contamination for the two samples (p>0.95).

We combined the genomic data from the two West Mound neonates with published East Mound Çatalhöyük genomes (Yaka, Mapelli, et al. 2021) as well as 195 published ancient genomes from early Holocene (Epipaleolithic to Chalcolithic) Southwest Asia, including both shotgun sequenced and SNP capture-based data. A timeline of the genomes used is presented in Figure 1.

**Figure 1.**
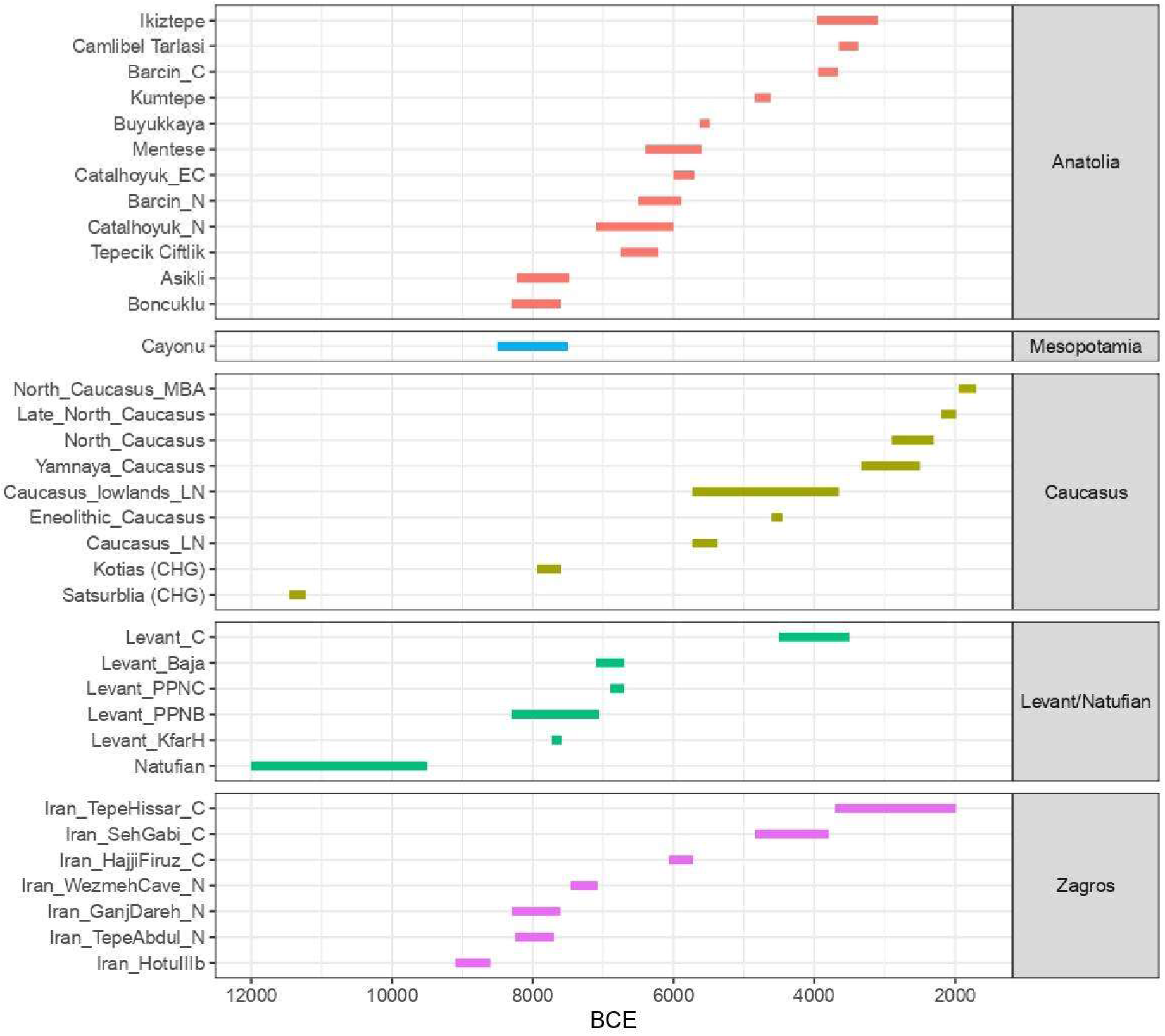
Timeline of ancient Southwest Asian and European individuals included in this study. The dates of the samples have been collected from the literature [Allen Ancient DNA Resource (AADR) (https://reich.hms.harvard.edu/allen-ancient-dna-resource-aadr-downloadable-genotypes-present-day-and-ancient-dna-data); v52.2] and include both direct C14 dates and stratigraphy-based dates.

### Genetic kinship analysis

We studied genetic kinship between the two West Mound neonates, which presents an interesting question given the low frequency of co-buried close biological relatives in published Çatalhöyük East Mound data (Yaka, Mapelli, et al. 2021). We estimated genetic kinship coefficients (θ) between the two West Mound neonates and also across all published genomes of East Mound Çatalhöyük. We used two different software, READ (Kuhn et al. 2018) and ngsRelate v.2 (Hanghøj et al. 2019). Only pairs with >5000 overlapping SNPs were included. Pairwise mismatch rates (P0) were calculated by READ, and normalized using the median P0 of all comparisons of Çatalhöyük individuals (i.e. assuming the average pair is not closely related, which is reasonable given the diversity of the sample in time). Genetic kinship coefficients (θ) were calculated as (1 - normalized P0). Meanwhile, ngsRelate was used to estimate θ using background population allele frequencies, for which we used all published Neolithic Anatolian populations as background. These methods, with some degree of uncertainty, can estimate kinship down to second (e.g. avuncular) or third degree (e.g. first cousin).

The results are shown in Figure 2. We replicated the earlier observation of a likely first degree relationship of a co-buried Çatalhöyük pair in Building 50 of the East Mound (Yaka, Mapelli, et al. 2021). Two West Mound neonates show no close kin relation with East Mound individuals, which is expected as the latest East Mound individual in our sample was dated to 6235–6075 (95%) (Düring and Marciniak 2005) (#Rosenstock et al. Chronology#), and is thus likely at least a century older than the West Mound neonates. The West Mound neonates also revealed no close genetic relationship between themselves (see Discussion).

**Figure 2.**
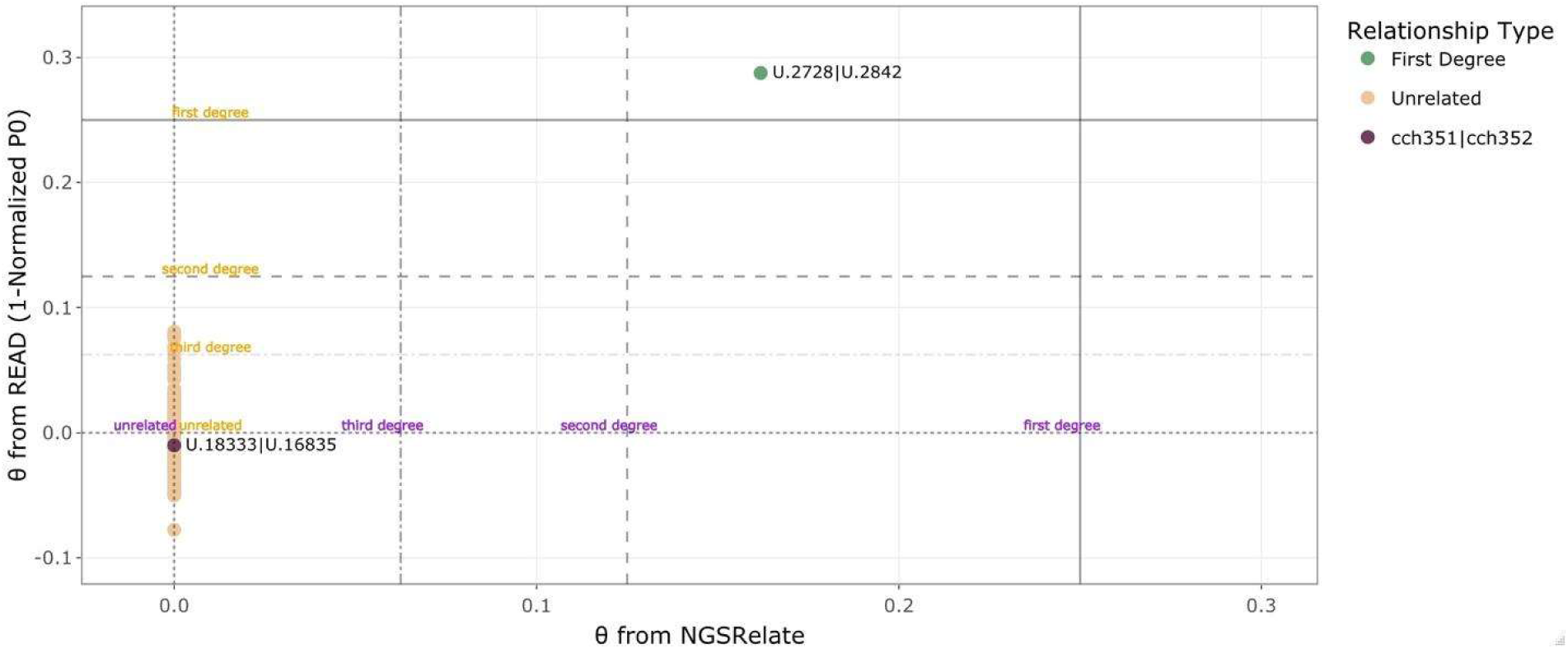
Kinship analysis results of Çatalhöyük genomes. Each dot represents a pair of individuals, which include the two West Mound neonates and 13 published genomes of East Mound Çatalhöyük (Yaka et al., 2021). The x-axis shows the genetic kinship coefficient (θ) calculated using the ngsRelate v.2 (Hanghoj et al., 2019) software, while the y-axis shows θ calculated using READ (Kuhn et al., 2018). The expected values (average θ) for first-, second-, third-degree related and unrelated pairs are marked as horizontal and vertical lines. The green dot shows the pair U.2728 - U.2842, from B.50 of the East Mound (identified in Yaka et al., 2021), and the purple dot shows the West Mound neonate pair.

### Population genetic characterization of the West Mound neonates

We next investigated population genetic affinities of the two West Mound neonates in comparison with genomes from Ceramic/Pottery Neolithic and Chalcolithic sites in Anatolia, as well as early Holocene groups from Southwest Asia (Figure 1). To obtain an initial overview, we carried out principal components analysis (PCA), using modern-day west Eurasian populations to calculate principal components and projecting ancient genomes onto the principal component space using the smartpca program of EIGENSOFT (Patterson et al. 2006). All Anatolian Pottery Neolithic (PN) individuals, including genomes from Central Anatolia PN [Çatalhöyük East Mound (Çatalhöyük_N) and Tepecik-Çiftlik] and from West Anatolia PN [Barcın (Barcın_N) and Menteşe] were clustered together in PC space, in close proximity to Aceramic Neolithic Central Anatolian groups (Aşıklı and Boncuklu), and largely distinct from roughly contemporaneous groups of Upper Tigris (Çayönü), South Levant (‘Ain Ghazal), or Central Zagros (Ganj Dareh) (Figure 2). Meanwhile, genomes from Chalcolithic sites from Anatolia, starting with a 6th millennium genome from Büyükkaya and following with the 4th millennium genomes from İkiztepe, Barcın, Kumtepe, and Çamlıbel Tarlası, showed a shift in PC space toward the Caucasus hunter-gatherers (CHG), in line with the inferred post-Neolithic “eastern” migration into Anatolia (Altınışık et al. 2022; Kılınç et al. 2016; Lazaridis et al. 2016; Skourtanioti et al. 2020) (Koptekin et al., under revision; also see Introduction).

Interestingly, the two West Mound neonates also fell clearly within the Central Anatolian Neolithic gene pool within the PC space, with no indication of “eastern” admixture observed in the Büyükkaya genome (Figure 3). We then tested these patterns using D-statistics using the qpDstat in AdmixTools software (Patterson et al. 2012). We computed D-statistics of the form D(*Outgroup, test; pop1, pop2*). The 1000 Genomes dataset Yoruba sample was used as an outgroup. Here, basically, if D < 0 this reflects higher genetic affinity between the *test* sample and the *pop1* sample, and if D > 0 there is higher genetic affinity between *test* and *pop2*. We used absolute Z score >2 as a measure of nominal statistical significance (not corrected for multiple testing).

**Figure 3.**
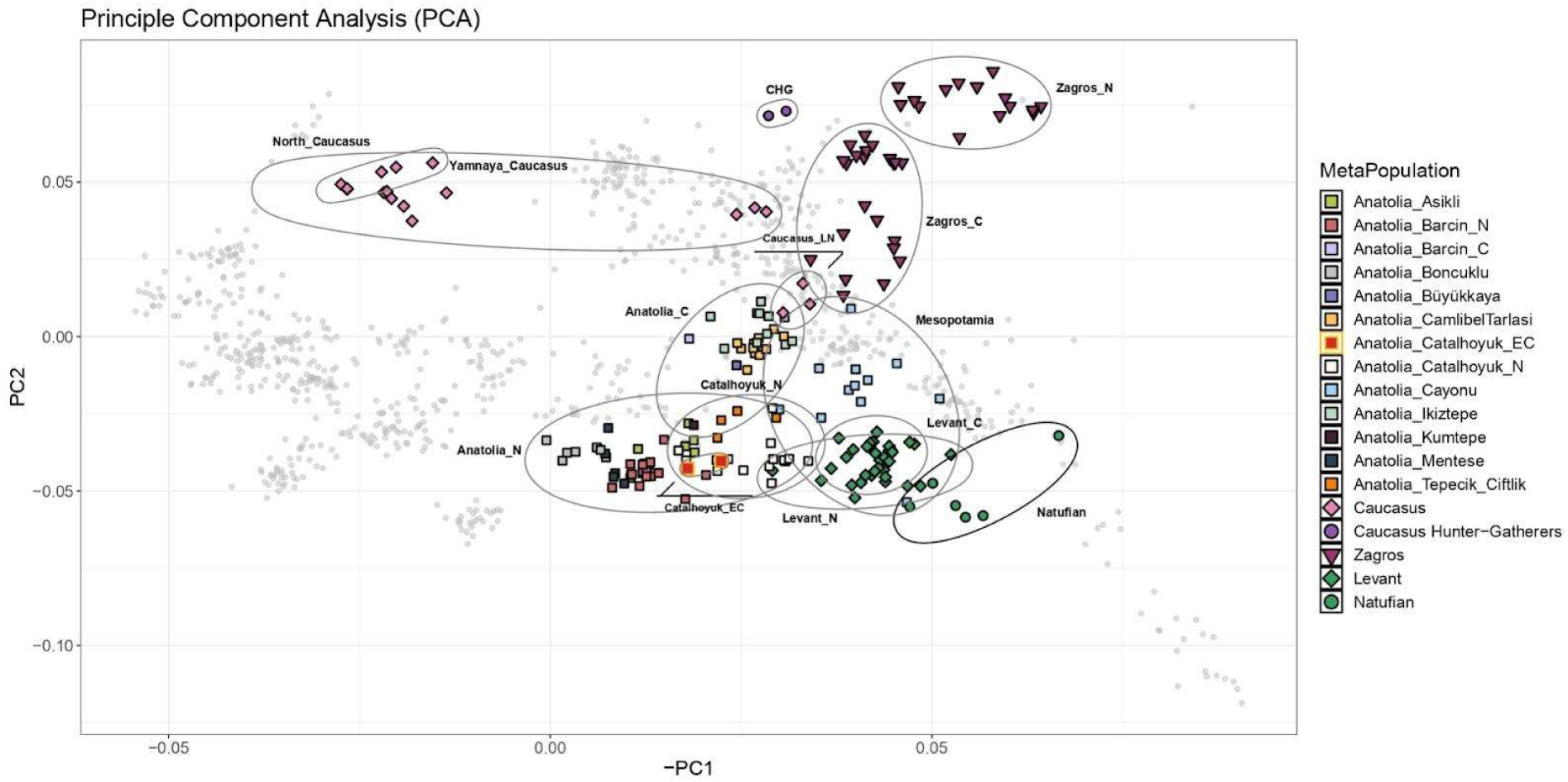
Principle Component Analysis (PCA) of Southwest Asian and European early Holocene populations. The PCs were calculated using 2068 individuals from 55 present-day West Eurasian populations (grey small dots) and 209 ancient genomes (colored symbols) were projected onto this space. Çatalhöyük Early Chalcolithic individuals fall into the Anatolian Neolithic cluster together with Çatalhöyük Neolithic individuals. Anatolia_N, Anatolia_C, Catalhoyuk_N, Catalhoyuk_EC, Zagros_N, Zagros_C, CHG, Levant_N and Levant_C stand for Neolithic Anatolian populations (Barcın, Çatalhöyük, Menteşe, Tepecik Çiftlik, and Çayönü), Chalcolithic Anatolian populations (Barcın, Büyükkaya, Çamlıbel Tarlası, İkiztepe, and Kumtepe), Neolithic Çatalhöyük, Early Chalcolithic Çatalhöyük, Neolithic Iran, Chalcolithic Iran, Caucasus Hunter-Gatherers, Neolithic Levant, and Chalcolithic Levant populations, respectively.

These D-test results revealed a number of observations. First, the Çatalhöyük West Mound neonates showed non-significantly higher affinity to each other than to any other Çatalhöyük (East Mound) genomes (Z>1.6). The Çatalhöyük West Mound genomes also tended to show higher affinity to the published 13 Çatalhöyük East Mound genomes (Catalhoyuk_N) than to other Anatolian Neolithic or to Anatolian Chalcolithic genome samples, albeit non-significantly (data not shown).

Second, we observed that eastern populations (e.g. Caucasus, Iran_N) showed higher affinity to most Chalcolithic Anatolian genomes (Büyükkaya, Ikiztepe, Çamlıbel Tarlası) than to Neolithic Anatolian genomes (Catalhoyuk_N, Tepecik-Çiftlik, Barcin_N, Menteşe) (Figure 4). This is in line with earlier observations on eastern gene flow into Anatolia, and with the patterns deduced from the PCA (Figure 3). Meanwhile, eastern populations showed similarly higher affinity to Chalcolithic Anatolian genomes than to Çatalhöyük West Mound (Catalhoyuk_C) (Figure 5), and showed no affinity towards Çatalhöyük West Mound over Neolithic Anatolia (Figure 6). Neither could we identify any neighboring population (Caucasus, Iran, or Levant) showing higher affinity to Çatalhöyük West Mound over East Mound (Figure 6). These observations overall imply the lack of any major gene flow into Çatalhöyük from a genetically distant source (e.g. CHG-related populations) by the turn of the 6th millennium BCE, that is by the time the East Mound was abandoned and the West Mound was established.

**Figure 4.**
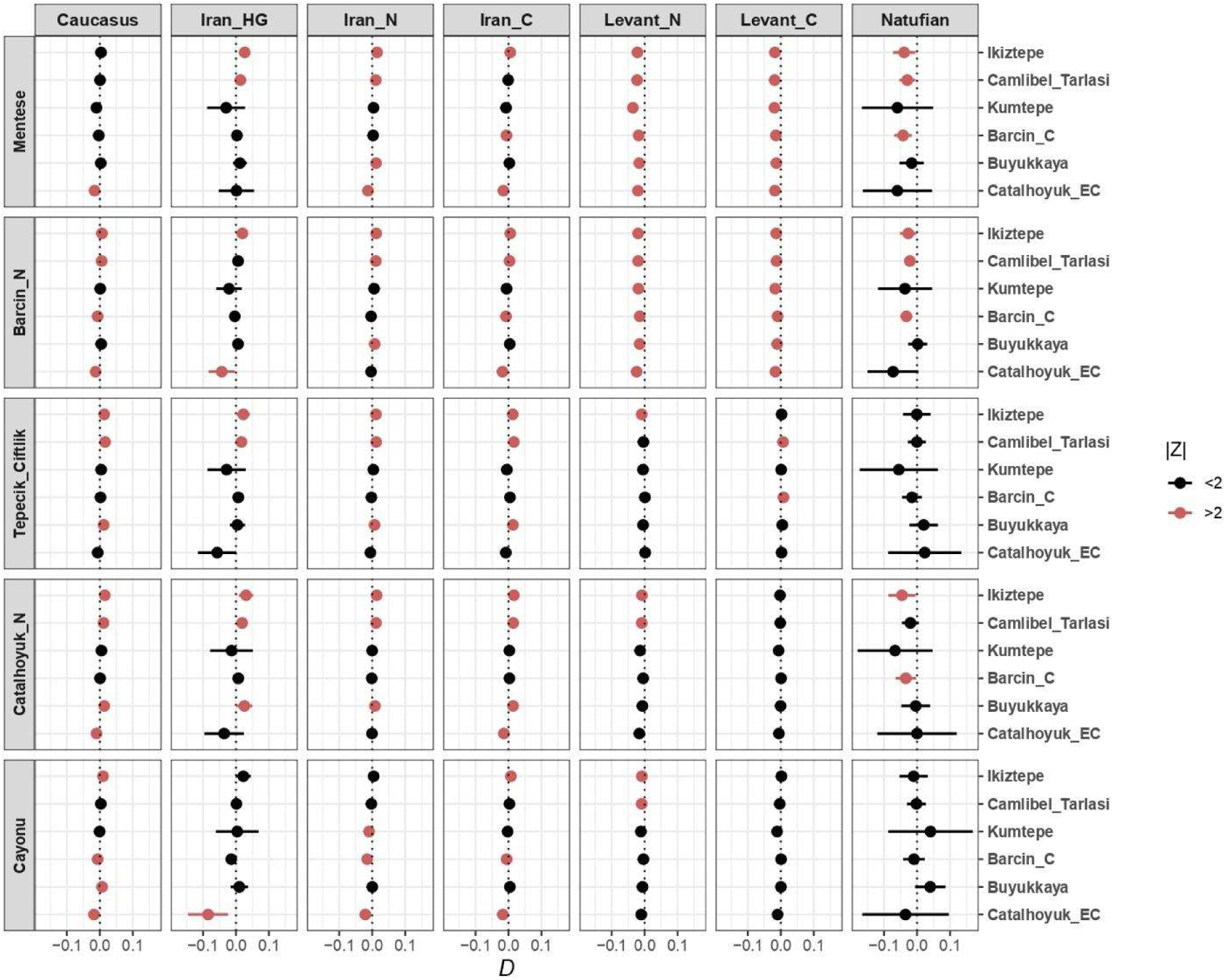
Genetic affinities of Çatalhöyük Early Chalcolithic individuals and Çatalhöyük Neolithic individuals with neighboring Anatolian populations. Results of D-statistics in the form of D(Yoruba, pop1; Anatolia_N, Anatolia_C). Horizontal bars represent ±2 standard errors. Catalhoyuk_N, Catalhoyuk_EC, Barcın_N, Barcın_C, Iran_HG, Iran_N, Iran_C, Levant_N and Levant_C stand for Neolithic Çatalhöyük, Early Chalcolithic Çatalhöyük, Neolithic Barcın, Chalcolithic Barcın, Iran Hunter-Gatherers, Neolithic Iran, Chalcolithic Iran, Neolithic Levant, and Chalcolithic Levant populations, respectively.

**Figure 5.**
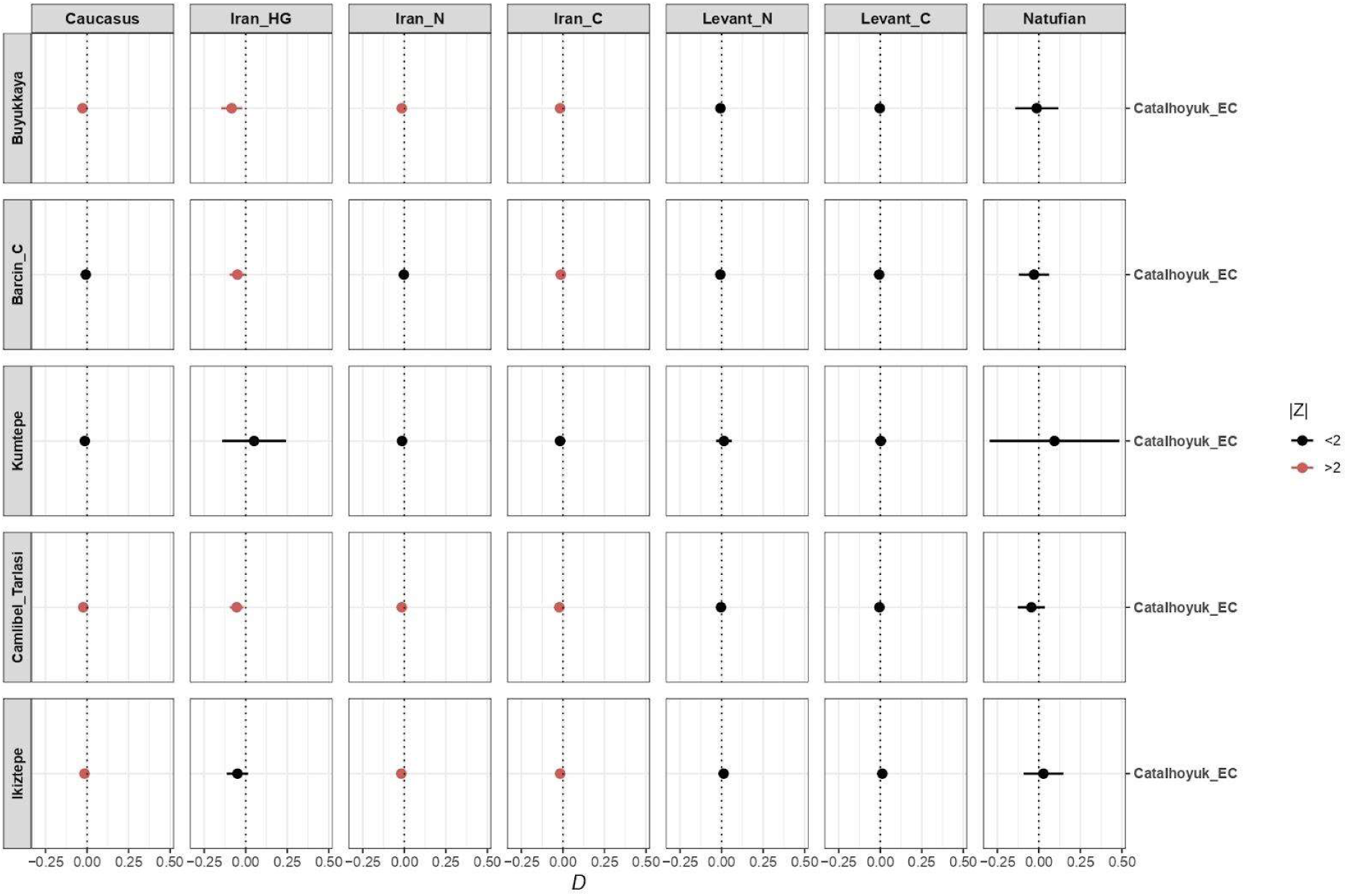
Genetic affinities of Çatalhöyük Early Chalcolithic individuals and neighboring populations. Results of D- statistics in the form of D(Yoruba, pop1; Anatolia_C, Çatalhöyük_EC). Horizontal bars represent ±2 standard errors. Catalhoyuk_EC, Barcın_C, Iran_HG, Iran_N, Iran_C, Levant_N and Levant_C stand for Early Chalcolithic Çatalhöyük, Chalcolithic Barcın, Iran Hunter-Gatherers, Neolithic Iran, Chalcolithic Iran, Neolithic Levant, and Chalcolithic Levant populations, respectively.

**Figure 6.**
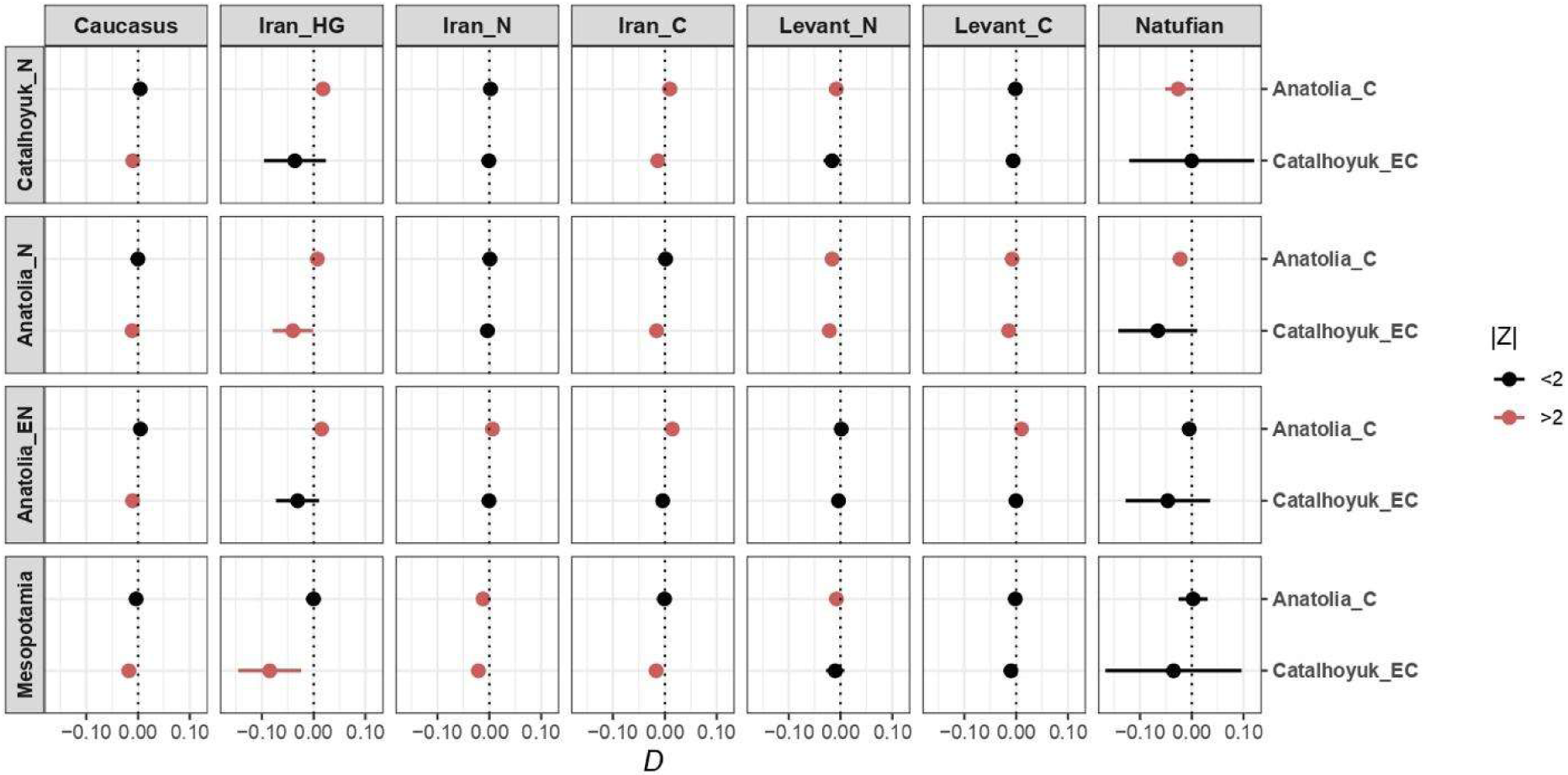
Genetic affinities of Çatalhöyük Early Chalcolithic individuals and neighboring populations. Results of D- statistics in the form of D(Yoruba, pop1; Anatolia_N, Anatolia_C). Horizontal bars represent ±2 standard errors. Catalhoyuk_N, Catalhoyuk_EC, Anatolia_EN, Anatolia_N, Anatolia_C, Iran_HG, Iran_N, Iran_C, Levant_N and Levant_C stand for Neolithic Çatalhöyük, Early Chalcolithic Çatalhöyük, Early Neolithic Anatolian populations (Aşıklı and Boncuklu), Neolithic Anatolian populations (Barcın, Çatalhöyük, Menteşe, and Tepecik Çiftlik), Chalcolithic Anatolian populations (Barcın, Büyükkaya, Çamlıbel Tarlası, İkiztepe, and Kumtepe), Iran Hunter-Gatherers, Neolithic Iran, Chalcolithic Iran, Neolithic Levant, and Chalcolithic Levant populations respectively. Mesopotamia includes Cayonu samples.

## Discussion

Here we report preliminary analyses of partial genomes from the only two Chalcolithic individuals yet recovered from Çatalhöyük West Mound. The data available is minimal, but it does allow inference on close genetic kinship and overall population genetic dynamics. We find that the two neonates, found in the infill of the same building, were not close kin, at least down to first cousins. This is consistent with earlier described patterns of frequent absence of close kinship among co-burials during the preceding East Mound at Çatalhöyük (Yaka, Mapelli, et al. 2021) (see Introduction). Nevertheless, we cannot reach any generalisation regarding burial traditions based on this single pair, especially given that the intramural burial tradition of also adult individuals in East Mound Çatalhöyük has clearly been abandoned for the West Mound (Brady et al. 2022) (Byrnes and Anvari, this volume).

Second, we find no evidence for a major admixture event that changed the Çatalhöyük gene pool between the end of the East Mound and the start of the West Mound. Although our genome coverage is limited, had there been an “eastern” gene flow as reflected in the Büyükkaya genome, we would have detected such signal. The Büyükkaya evidence is based on a single genome dated to 5626-5515 cal BCE (Skourtanioti et al. 2020), but the “eastern” influence in the Anatolian post-Neolithic gene pool is also supported by unpublished genomes from our group dating to the mid- 6th millennium BCE. The fact that we do not observe this signature in Çatalhöyük West Mound by 5800 BCE can be explained by two scenarios: either this “eastern” admixture event initiated approximately between 5800 BCE and 5600 BCE, or it started earlier but Çatalhöyük remained isolated. Further data from Anatolian sites dating to the very early 6the millennium BCE can help directly resolve this question. Whether the source of the admixture could be related to the populations from the Caucasus/northeast Anatolia, or populations associated with the north Mesopotamian Halaf culture (#Rosenstock, the developed Neolithic#), and whether it may be related to the hiatus observed in Anatolian settlements by the mid-6th millennium, may also be eventually resolved by denser genetic sampling.

For now, we can conclude that the changes in subsistence and social organization between Çatalhöyük East and West Mound were not impacted by any large-scale human movement from distant areas with distinct genetic profiles. We note, however, that we are here assuming that the two neonates were not genetic outliers with distinct ancestry from the rest of the West Mound population. This issue, in turn, could only be resolved by recovery of other burials from the site.

## Acknowledgements

We thank Eva Rosenstock, Jana Anvari, David Orton, all colleagues at the METU CompEvo, Hacettepe Human_G and Center for Palaeogenetics (CPG) for their support, suggestions and/or comments, and Arda Sevkar and Ayça Aydoğan for help in computational analyses. We also thank the Konya Museum and the Ministry of Culture for permissions to work on the materials. The authors acknowledge support from H2020 ERC Consolidator Grant (no. 772390 “NEOGENE” to M.S.); YOK 100/2000 Doctoral Scholarship; TUBITAK 2211/A Doctoral Scholarship, and Hacettepe University Scientific Research Projects Coordination Unit (no: 16769 and 14528 to A.M.B.). Computations were performed at NEOGENE (Middle East Technical University).

## Notes

### Competing Interest Statement

The authors have declared no competing interest.

